# The morphotype approach to classification of aerial animals in radar data

**DOI:** 10.64898/2026.03.24.713935

**Authors:** Yuval Werber

**Author notes:** Address: Department of Evolutionary and Environmental Biology, University of Haifa 199 Aba Khoushy Ave. P.O. Box 338, Haifa, 3103301, Israel. **Open research:** Data and code are provided as private-for-peer review via the following links: Data: https://doi.org/10.6084/m9.figshare.32944700, Code: https://doi.org/10.6084/m9.figshare.32944709.

## Abstract

Radar aeroecology makes ecological inferences about aerial wildlife using radars. While generating unique, large-scale datasets describing airborne biological activity, radars are largely incapable of relating detections to specific species and are thus taxonomically limited. I describe an analytical method to increase taxonomic resolution in vertical looking radar data by dividing detected organisms into morphology- and movement-based “aerial morphotypes”. The Birdscan MR1 radar, widely used in aerial wildlife research and monitoring, is deployed in dozens of locations across Europe, the Near East and the United States. Using the MR1’s target classifier, wing flapping frequency calculations and target size estimations, I demonstrate a nearly 8-fold increase in bird classification resolution using radar data from the Hula Valley Research station, Israel. I show that most newly defined morphotypes correspond to small numbers of local species (1-10). Thus, the approach provides a realistic opportunity to bridge the taxonomy gap in radar aeroecology. By using the approach, radar aeroecologists can estimate “Aerodiversity”, analogous to biodiversity, a fundamental, widely used measure in ecology and conservation. By explicitly addressing taxonomy in radar aeroecology, practitioners will increase the impact and dissemination of their work and contribute to a deeper, more complete understanding of the aerial habitat.

## Introduction

The airspace is an inseparable part of life on earth (Diehl 2013, Diehl et al. 2017). Trillions of organisms, from bacteria, through plants and fungi to insects and various sized vertebrates use it daily for mobility, foraging, and other behaviours (Cain et al. 2000, Finlay 2002, Huang et al. 2024). From a scientific perspective, the ecological importance of aerial habitats is contrasted by the limited ability to monitor and study them, as they are largely out of human reach, complicating direct observations (Bauer et al. 2019). Radar technology is the main tool presently available to conduct large-scale observations of biological activity in the airspace. The field of radar aeroecology has evolved along with hardware and data processing technologies, enabling far-reaching inferences related to behavioral ecology, macro-ecology, biogeography, and conservation biology.

Vertical looking radars (VLR) are designed to conduct individual-level monitoring in small air volumes (∼1 km^3^). VLRs identify, track, and characterise single biological targets, from minute insects to pelicans, at ranges far greater than what is humanly possible to observe (Chilson et al. 2017). The automated operation of modern VLRs enables long term, consecutive monitoring, efficient data processing, and large-scale analysis. In addition to their role in aeroecological research, field experiments, and long term monitoring of aerial habitats (Nilsson et al. 2018, Werber et al. 2023a, Knop et al. 2023, Weisshaupt et al. 2023, Haest et al. 2025, Werber and Sapir 2025, Jimenez et al. 2025, Werber et al. 2025), VLRs are also used for aviation safety and wildlife conservation in the vicinity of wind farms and other tall infrastructure that protrude into the airspace (Köppel 2017, Werber 2024). In this capacity, VLRs provide unmatched individual level monitoring capabilities in terms of temporal resolution and spatial accuracy.

VLRs work by emitting radio waves and analysing the echoes returned from bodies of airborne animals - specifically the water content in their tissues. The resulting data consist of precisely timed (to the 1000th of a second) echo intensity-time sequences lasting up to dozens of seconds (depending on flight altitude, track and target size), during which detected animals may fly over hundreds of meters (Schmid et al. 2019). VLR algorithms calculate descriptive parameters related to target size, shape, and movement patterns (Chapman et al. 2011, Feng, H. 2025). Wing flapping patterns, central to VLR-based research, can be accurately resolved and contain taxonomic “hints”, as these patterns correspond to animal taxonomy and organism size (Schaefer 1976, Able 1977, Pennycuick 2001, Zaugg et al. 2008, Werber et al. 2023b). Wing flapping constitutes an axis between biomechanics, taxonomy, behavior and energetics with immense research potential, especially in big-data frameworks applied in radar-based ecological research.

Currently, the main limitation of radar-aeroecology is the inability to directly discern the taxonomic identities of nearly all detected organisms. While emerging AI approaches do allow coarse taxonomic classification (Zaugg et al. 2008, Werber et al. 2023b), the resolution lags far behind standard taxonomic accuracy in other fields. Taxonomic “blindness” inhibits radar aeroecology and the dissemination of its methodologies and scientific output. I present an analytic approach intended to bridge a large part of the radar-taxonomy gap by further resolving coarse radar-produced classes into finely-separated “aerial morphotypes”. These can act as operational, non-taxonomically specific, functionally meaningful units that are better correlated with local species inventories (Figure 1). The approach drastically reduces taxonomic uncertainty in VLR data, in many cases narrowing down candidate taxa to a single species or a few potential species at a given time and place (Figure 3). Applying the approach to rapidly growing VLR datasets can improve inference regarding Aerodiversity (biodiversity’s airspace equivalent) and associated key ecological processes, open new avenues of research, and streamline the integration of radar aeroecology with other scientific fields.

**Figure 1:**
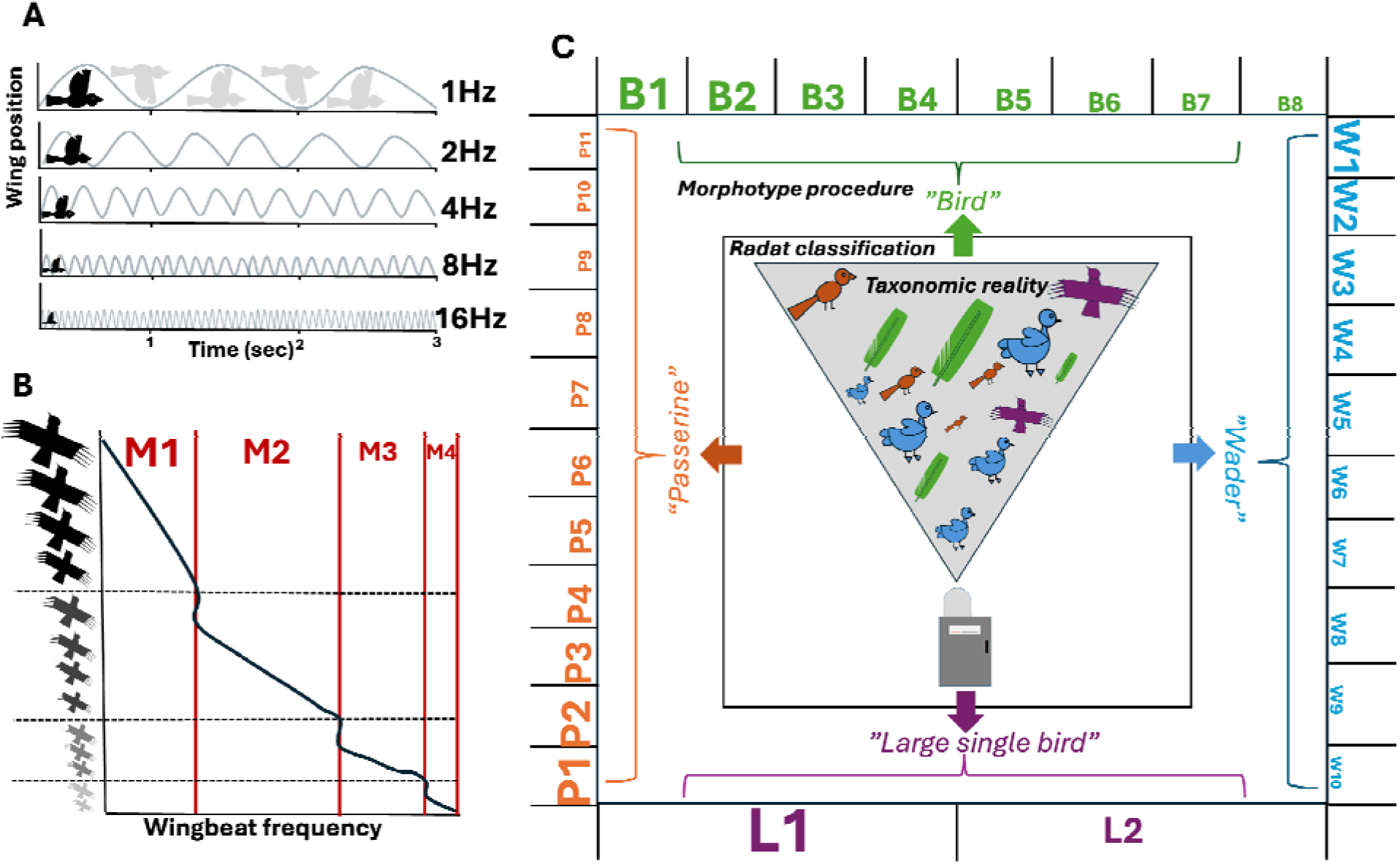
Method summary. A. Schematic description of Wing flapping frequency (WFF) and its negative relationship with body size. Grey curves signify changes in body position during flapping flight, with each wave representing a single flapping cycle. B. Schematic description of the morphotype separation procedure based on WFF-body size relationship. Red lines and different bird shading signify morphotype separation based on sharp leaps in WFF between adjacent size groups. M is short for “Morphotype”. C. Schematic description of complete morphotype procedure: Grey triangle represents the actual occurrence of individual birds from different species in the radars’ detection range, having different sizes and taxonomic/morphologic affiliation. Inner square signifies the embedded MR1 classifier, lumping individuals to one of four classes. Outer square represents the morphotype procedure, applied over the MR1 classifier to bin classified target into morphologically and functionally meaningful groups. All icons and graphics created by the author, Yuval Werber.

**Figure 2:**
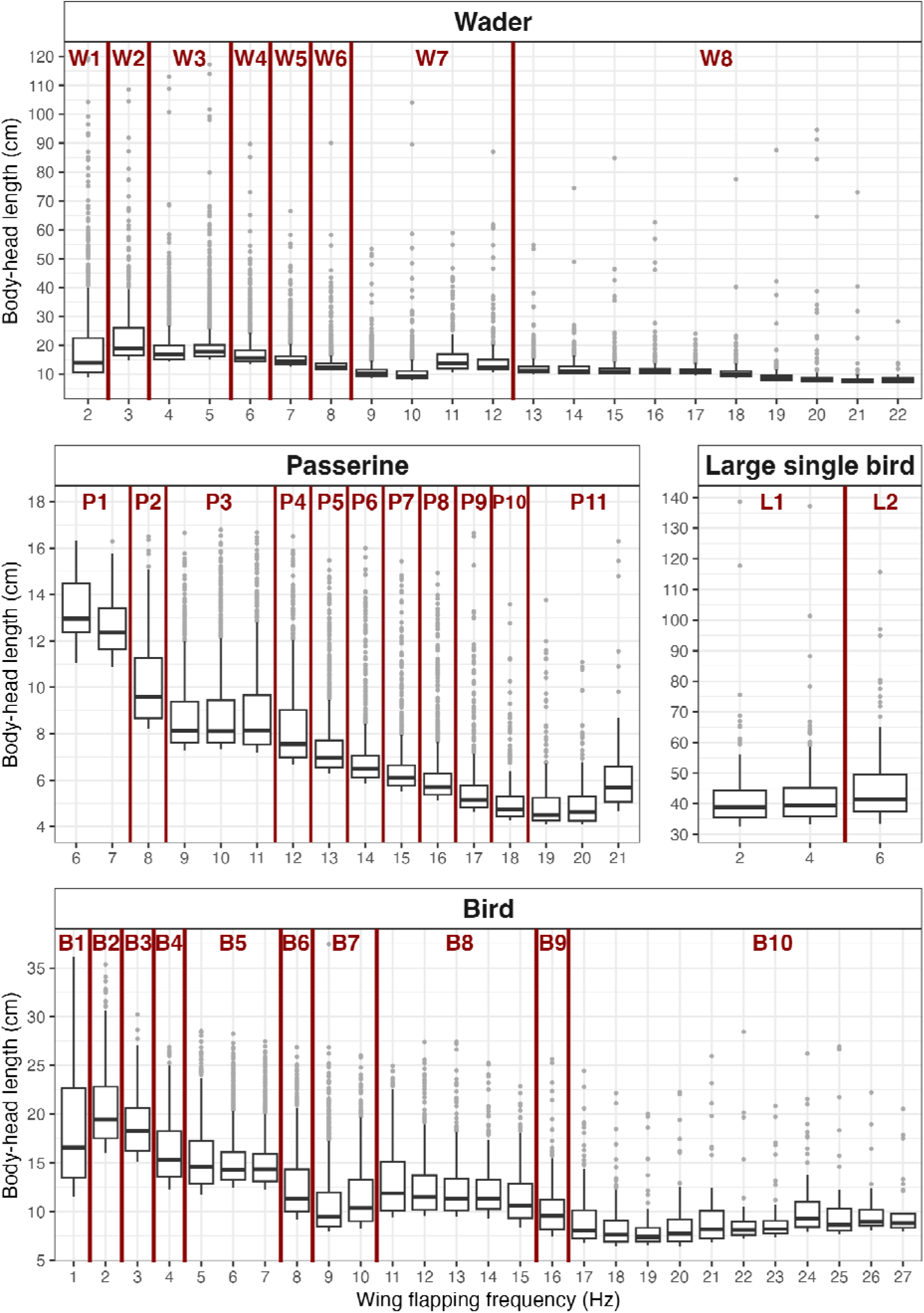
Resulting morphotypes following segregation procedures. Each panel describes the results of a different radar class. Boxes represent 1 or 2 Hz WFF intervals and red lines illustrate significant segregation between adjacent WFF intervals, used to define segregated morphotypes. Within the boxes, the middle horizontal lines represent median values, edges represent upper and lower quartiles, whiskers represent ±1.5*interquartile range.

**Figure 3:**
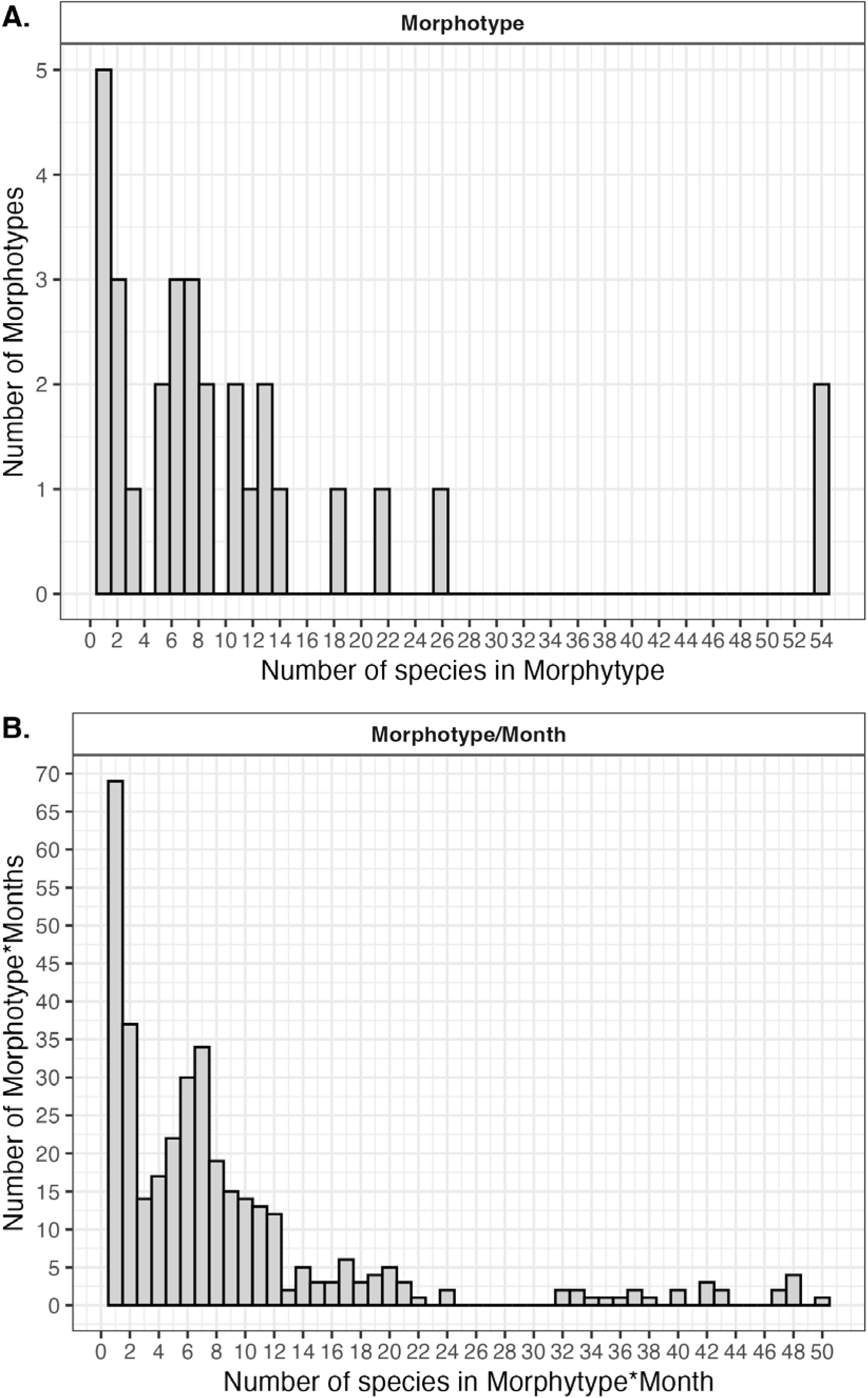
A. Histogram of the number of bird species comprising each morphotype. B. Histogram of the number of species comprising each morphotype/month.

### Case study

I applied the morphotype approach to a BirdScan MR1 VLR database collected in the Hula Valley, Israel, and used the regions’ species inventory to explore the method’s impact in a complete taxonomic context. The Valley is a major stopover site for huge populations of hundreds of migratory species, which breed across the palearctic from Western Europe to Siberia and funnel through the bottleneck between the Arabian Desert and Mediterranean Sea (Shirihai et al. 1996, Leshem and Yom-Tov 2008). Millions of Migrants stop along the waterways and productive habitats in the region to refuel ahead of crossing the Sahara or landing to recover once the crossing is complete. Given its location and spatial circumstances, the Hula Valley is a hotspot of taxonomic and behavioral diversity featuring a wide spectrum of morphologies and phenologies. Mild winters and consistent water supply from natural springs mean the region remains highly productive year-round, maintaining high terrestrial and aerial biodiversity (Rotem and Weil 2014).

The radar station in the Hula Valley Research Center, Israel (35°430 E, 33°030 N), which has been in continuous operation since August 2018, is the longest continuously operated station of the Israeli BirdScan network consisting of 11 stations spread across the country. Its location in a major migration stopover area makes it ideal for exploring radar data from a diverse aerial community with substantial turnover throughout the year (Shirihai et al. 1996, Shy, E., Beckerman, S., Oron, T. and Frankenbergg, E., 1998, Gophen 2023). The region is also a global birding hub, with armatures and professionals monitoring it’s avian fauna for several decades (Collins-Kreiner et al. 2013). This unique mix of ecological circumstances, long term monitoring and radar deployment form an optimal setup to assess the method’s practical impact. I use these resources to obtain two measures of efficiency for the morphotype method. First, I assess the level of increase in “taxonomic resolution”, meaning the increase in class separation within the radar data. Second, I project this increased resolution onto species-level taxonomic reality by associating each species in the region’s inventory to its relevant morphotype, and explore the extent to which the method improves our observation capability.

## Methods

### Data collection and preparation

The BirdScan MR1 (Swiss BirdRadar Solution AG, https://swiss-birdradar.com/systems/radar-birdscan-mr1/, wavelength 3.2 cm, 9.4 GHz, 25 kW, software version 1.6) is an X-band VLR with a wide aperture (∼60°), 20 dB horn antenna. Its random forest-based classifier automatically distinguishes birds (“Wader”, “Passerine”, “Large single bird”, “Bird”), insects, flocks, non-biological targets, and recently also bats (Zaugg et al. 2008, Werber et al. 2023b). The radar can detect the smallest flying vertebrates (Chiffchaff, *Phylloscopus collybitaI*; Common pipistrelle, *Pipistrellus pipistrellus)* up to ∼1,000 m above ground level (AGL), while larger species are detectable at much higher altitudes.

The morphotype approach is demonstrated using a bird dataset recorded by the Hula Valley BirdScan radar between August 2018 and March 2021. I only used data collected in the “Short pulse” operation mode (pulse length 65 ns, pulse rate frequency 1800 Hz, range resolution ∼7.5 m), optimal for individual vertebrate detection; which was used for two-thirds of every hour during the data collection period. Data from rainy periods, when the radar’s output is not reliable, were omitted by coupling each detection with its closest rain measurement (to within 10 minutes) from a nearby meteorological station and removing those with more than 0 mm of rainfall. Detections recorded below 50 m AGL were removed due to the prevalence of ground clutter in data from these altitudes, and detections from above 1,000 m AGL were removed to maintain detectability of the full taxonomic range of birds in the region. I only retained detections classified as one of the bird classes (“Wader”, “Passerine”, “Large single bird”, “Bird”) with a classification probability (automatically outputted by the radar’s classifier) higher than 50%. Insect detections were excluded based on their classification by the BirdScan’s classifier and bat detections were removed using the BATScan classifier according to Werber et al. (2023b).

### Morphotype segregation

The morphotype segregation process is based on the widespread observation that wing flapping patterns among all taxa of flying animals are related to size and taxonomy (Schaefer 1976, Pennycuick 1990, Bruderer and Popa-Lisseanu 2005, Bruderer et al. 2010). Wing flapping frequency (WFF, Hz) and radar cross section (RCS, cm^2^) are both automatically calculated for each detection by the radar. WFF, the number of wingbeats the organism flaps per second during flight (Figure 1A), is automatically calculated by the radar based on power oscillations in the raw echo (echo intensity, dB, over time), which are caused by changes in body shape and posture relative to the radar as the animal flaps its wings (Zaugg et al. 2008, Gong et al. 2022). WFF is determined as the dominant frequency component in a fast Fourier transformation of the raw signal, automatically executed by the BirdScan radar (see Werber et al. 2023b for a visual depiction).

RCS is a measure of the amount of radiation reflected back from the detected organism to the radar’s antenna, and in optimal detection conditions, it is descriptive of animal size as downward facing, wet tissue surface area (Schmid et al. 2019, Schmaljohann 2020). The main reflective substance in biological targets is water, while dry tissue (chitin, hair, feathers) does not have any effect on the RCS (Bruderer 1997, Chilson et al. 2018). For birds, whose wing surface and tail are largely made of feathers, RCS describes the bottom surface area of the body and head, as documented from below by the radar (Chilson et al. 2018). In vertical looking radars, RCS is highly sensitive to the manner by which the organism crosses the detection range and is regarded as an accurate estimation of organism size (as body surface area) only when the animal flies parallel to the ground (no pitch or bank) directly above the radar. Accuracy of estimations increasingly declines for targets crossing towards the periphery of the detection area and those flying in pitched or banked postures. Importantly, calculated RCS cannot be larger than the targets’ actual bottom surface area because an object can only return a finite amount of radiation. Crossing through the periphery of the detection area and in pitched or banked postures can only reduce the amount of radiation reflected back to the radar by either reducing the radiation initially projected on the animal from the radar or shrinking the surface area exposed to radiation, respectively. Thus, the measure can only be biased towards lower RCS values, meaning that for a given group of signals derived from identical organisms, the highest measured RCS value will be the most accurate one (Schmid et al. 2019).

I rely on these notions to separate the main bird classes produced by the radar’s classifier into subgroups of distinct wing flapping frequencies and statistically significant size differences, creating groups with unique, non-overlapping wing flapping frequency-radar cross section (WFF-RCS) combinations (Figure 1A,B). The raw data is split into WFF integer interval bins (henceforth WFF intervals), each representing a potentially morphologically-functionally distinct group, and the RCS values in each bin are compared with adjacent bins to validate or reject the distinction. This involves using a small subset of the data (5% in the presented procedure) in which size estimation is accurate to identify significantly distinct WFF intervals in each radar class and then extrapolating this grouping to the entire dataset based on the accurately measured WFF of each detection.

### Extracting accurate RCS subsets

A subset of radar targets with accurately estimated organism sizes is required for correct segregation into morphotypes. This subset was obtained by splitting the data from each radar class into 1 (“Wader”, “Passerine”, “Bird” classes) or 2 (“Large single bird” class) Hz WFF intervals and retaining only the top 5% of RCS values in each interval, according to Schmid et al. (2019). Given that organisms of the same MR1-produced radar class and WFF interval are morphologically similar (Schaefer 1976, Pennycuick 1990) and that RCS can only be biased downwards, and assuming the WFF calculation is correct, the largest 5% is expected to have an accurate RCS estimation, representing organism bottom surface area for the given WFF interval (Schmid et al. 2019). Body-head length was estimated by calculating the diameter of a circle with an area equal to the target RCS (cm^2^) using basic geometry (Werber et al. 2023b). For each original radar class, WFF intervals with fewer than 30 detections were aggregated to the larger adjacent WFF interval group, either below or above.

### Inference through multiple comparisons

The resulting dataset, containing only accurately sized detections (5% of all detections), was subject to a series of multiple comparisons in which the mean RCS of each WFF interval was compared to that of the ones just above and below it. When two adjacent WFF intervals had significantly different RCSs, they were regarded as representing birds of different morphotypes (Figure 1B). Significantly segregated morphotypes were named with the first letter of the parent radar class (“W”, for Waders, “P” for Passerines, “L” for Large single bird and “B” for Bird) and a serial number starting with one and continuing in ascending order, with lower numbers indicating slower WFFs (e.g. larger bodied morphotypes, Figure 1C). The morphotype segregation procedure was applied on the entire accurately sized bird detection dataset, resulting in a set of distinct morphotypes for each radar class (Figure 1C). The series of multiple comparisons was conducted by first applying an analysis of variance (ANOVA) using the “aov” function and then applying the “TukeyHSD” function on the result, both from base R. All processing, analysis and visualization were done in R Studio version 4.5.0.

### Hula Valley case study - Taxonomic survey

I used the “Society for the Protection of Nature in Israel”’s (SPNI) database to compile a list of all bird species occurring in the Mediterranean habitat in Israel (332 species of 21 orders, Table S1). Species that were only observed a single time throughout the data collection period (several decades), or those whose occurrence is unlikely based on expert opinion, were not included in the dataset. For each species, its genus, class, mass, body length ranges, and monthly prevalence were manually registered from the species’ page on the SPNI’s dedicated bird website (https://www.birds.org.il/he, Table S1). The website is routinely updated with expert-validated bird observations across the country. When information regarding particular species was missing from this database, I obtained it from relevant scientific literature. Next, each species was assigned a WFF estimation from published literature (mostly Bruderer et al. 2010, see Table S1 for additional species-specific references). For species with no published WFF range, I calculated presumed WFF as WFF=4.18xMass^-0.264^ according to Rayner (1995). Each species observed in Northern Israel was assigned to one of the four bird classes automatically outputted by the radar. All wader species (order Charadriiformes, 65 species) to the “Wader” class, passerine species (order Passeriformes, 141 species) were assigned to the “Passerine” class, and the rest were divided so that species with a head-body length larger than 30 cm were designated as “Large single bird” (92 species) and the remainder as “Bird” (33 species). This division represent the a priori taxonomic resolution of bird detections by the MR1 radar before the application of the morphotype approach.

The species assignment procedure was conducted twice; once without reference to time of year (resulting in a table consisting of the 332 observed species and their assigned morphotypes, Table S2) and once divided by month. In the monthly table, the species pool associated with each month was assigned a Morphotype/Month label, resulting in a table with 3,124 entries (Table S3), each representing the occurrence of a single species in a given month. Species with WFF that were not represented in the radar data (8 passerine species and 12 “Large single bird” species, <1% of all species) were given a morphotype name with no number.

## Results

### Morphotype segregation

The 4 parent radar classes were segregated into a total of 31 morphotypes of significantly different body size and WFF (“Wader”: 8 morphotypes, F=194, p<0.0001, “Passerine”: 11 morphotypes, F=600, p<0.0001, Large single bird: 2 morphotypes, F=4.7, p=0.009, Bird: 10 morphotypes, F=285, p<0.0001, Tables 1, S1, Figures 1C, 2), increasing classification resolution by a factor of 7.75 (from 4 to 31 different classes). The relationship between WFF and RCS largely agreed with the convention that slower flappers are larger, with significant negative correlation in the ‘Wader’ (Correlation coefficient: -0.2, p<0.0001), ‘Passerine’ (Correlation coefficient: -0.63, p<0.0001), and ‘Bird’ (Correlation coefficient: -0.48, p<0.0001) classes.

**Table 1:** Morphotype segregation summary: A list of segregated morphotypes including Average wing flapping frequencies (WFF), body-head lengths, and radar cross sections (RCS).

| Radar Class | Morphotype | Average WFF (Hz) | Average Body-Head Length (cm) | Average RCS (cm<sup>2</sup>) |
| --- | --- | --- | --- | --- |
| Wader | W1 | 3.41 | 20.88 | 498.94 |
| Wader | W2 | 5.05 | 19.42 | 354.88 |
| Wader | W3 | 6.34 | 19.16 | 336.60 |
| Wader | W4 | 7.97 | 16.12 | 228.62 |
| Wader | W5 | 8.94 | 13.55 | 158.97 |
| Wader | W6 | 9.85 | 11.03 | 104.53 |
| Wader | W7 | 12.16 | 12.89 | 164.50 |
| Wader | W8 | 17.82 | 11.24 | 123.44 |
| Passerine | P1 | 7.05 | 13.24 | 139.33 |
| Passerine | P2 | 8.00 | 12.52 | 123.92 |
| Passerine | P3 | 10.23 | 9.18 | 69.17 |
| Passerine | P4 | 11.95 | 9.02 | 67.19 |
| Passerine | P5 | 12.97 | 8.36 | 57.95 |
| Passerine | P6 | 14.01 | 7.52 | 46.34 |
| Passerine | P7 | 15.01 | 6.84 | 38.02 |
| Passerine | P8 | 15.99 | 6.39 | 33.02 |
| Passerine | P9 | 16.91 | 6.08 | 30.21 |
| Passerine | P10 | 17.89 | 5.55 | 25.54 |
| Passerine | P11 | 19.74 | 5.22 | 23.09 |
| Large Single Bird | L1 | 3.06 | 42.04 | 1506.70 |
| Large Single Bird | L2 | 4.99 | 43.43 | 1588.28 |
| Bird | B1 | 2.19 | 18.65 | 304.12 |
| Bird | B2 | 2.76 | 20.69 | 349.45 |
| Bird | B3 | 4.07 | 18.94 | 290.31 |
| Bird | B4 | 5.08 | 16.28 | 217.52 |
| Bird | B5 | 7.01 | 15.23 | 188.69 |
| Bird | B6 | 8.92 | 12.70 | 137.23 |
| Bird | B7 | 10.41 | 11.41 | 114.18 |
| Bird | B8 | 14.25 | 12.18 | 123.79 |
| Bird | B9 | 16.92 | 10.17 | 86.79 |
| Bird | B10 | 21.01 | 9.17 | 74.37 |

### Local species inventory case study

Following the application of the morphotype segregation process, each species in the local species inventory was assigned to a morphotype based on its taxonomy, size, and WFF. The resulting species distributions (Fig. 3) represent substantial improvement compared to the a priori taxonomic resolution produced by the MR1’s classifier.

Regardless of the time of the year, 5 morphotypes (15%) included only a single species, 9 morphotypes (28%) included less than 5 species, and 19 morphotypes (59%) included less than 10 species (Table S2, Figure 3A). At a monthly resolution, 69 morphotype/months (19%) were attributed to a single species, 137 (38%) were attributed to less than 5 species, and 257 (71%) were attributed to less than 10 species (Table S3, Figure 3B). The number of morphotypes and morphotype/months attributed to a large number of species was generally low (Fig. 3 A,B).

## Discussion

The morphotype approach described in this paper leverages robust radar-derived parameters and simple statistics to improve taxonomic resolution in radar aeroecology. The case-study analysis of birds from the Hula Valley shows that the majority of morphotypes are associated with a narrow set of species, and in many cases only a single one. This supports the conjecture that radar methodologies can greatly benefit from taxonomic input and challenges the common practice of explicitly avoiding radar-based taxonomic assignment.

The approach can easily be applied to any BirdScan dataset using the published code and local species inventories, and is potentially applicable to any VLR that produces individual-level observations and outputs morphologically meaningful parameters. Bat fauna, morphologically diverse and composed of much fewer species, can also be analyzed using the same approach. For insects, given their extensive diversity, replacing species with higher taxonomic levels and incorporating body shape-related parameters can enable productive application. While RCS is a basic feature of radar data, wingbeat information may not always be available. Nonetheless, other body shape or movement-related parameters can be included in models, permitting extension of this approach to nearly any biological radar system (Weisshaupt et al. 2023, Huang et al. 2024).

While VLRs are better equipped to perform the described procedure compared to other systems (weather radar, wind profilers, etc.), there are other methods to relate radar data with taxonomic reality. Validation procedures using other information sources (cameras, acoustic recording, environmental DNA, LIDAR, citizen science, surveys) can substantially improve upon current approaches (Gauthreaux and Livingston 2006, Schmidt et al. 2017, Nilsson et al. 2018, Haas et al. 2022, Van Doren et al. 2023). Furthermore, as advances in AI increasingly integrate into scientific practice, target classification will also improve, providing opportunities to further bridge the radar-taxonomy gap. However, given that large-scale species-specific inference is an unlikely product of algorithmic and hardware advances in the near future, using morphotypes as morphology-based taxonomic proxies can create promising opportunities in research and conservation.

The use of morphotypes as proxies for groups of species, and in some cases, a single species, is biologically justified, especially in the aerial habitat. If animals differ in size and movement, their responses to, and interactions with the surrounding environment are also likely to differ (Pennycuick 2008), resulting in species- or morphotype-specific, physically derived niches (Nilsson et al. 2025). This is also often the case in entomology, botany, and mycology, fields in which species-specific identification in real time in the field is nearly impossible. Accordingly, higher taxonomic classifications based on morphology or functionality are used to describe diversity and make ecological inferences (Duckworth et al. 2000, Bellamy et al. 2018, Lofgren and Stajich 2021, Morel et al. 2022).

Radar aeroecologists routinely encounter concerns from scientists and practitioners outside the field related to the lack of taxonomic anchors in their data, which can challenge inference and translation. Yet, radar-derived scientific discoveries have fundamental, far-reaching implications, which are increasingly being recognized by the wider scientific community (Bauer et al. 2017, 2024, Shamoun-Baranes et al. 2019, Nathan et al. 2022). Bridging the gap between radar aeroecology and traditional taxonomy is a worthy undertaking that deserves attention and effort. The morphotype method presented here begins to close this gap, and its wider implementation will help to expand the scope, impact, and relevance of radar-based aeroecological research.

### Perspectives and applications

Increasing taxonomic resolution in radar data via the morphotype approach has scientific and applicative potential for novelty, as well as conservation significance. Scientifically, radar-derived morphotypes can enable the study of fundamental ecological concepts in the aerial habitat, extending our understanding of phenomena that have been well-researched in terrestrial and aquatic habitats for decades (Bloch et al. 2025). Richness and diversity measures, basic to our understanding and conservation of ecosystems (Whittaker et al. 2001, Brooks et al. 2006, Maclaurin and Sterelny 2008), can be introduced into radar aeroecology using functionally relevant morphotypes. By extending the application to bats and insects, the resulting observed diversity can facilitate the introduction of community ecology, which has contributed greatly to our understanding of nature, to aerial habitats (Gauch, H.G. 1982, Vellend 2010). Improved taxonomic resolution can also facilitate the exploration of particular cases (fine taxonomic or functional groups, specific behavioral phenomena and physiological processes) using radar data, opening new avenues for scientific use and ecological inference.

Radars are increasingly used to conduct preliminary surveys and subsequent monitoring of airspace around airports, wind farms and high structures. Increased functional resolution in radar products will improve prediction accuracy and threat identification, which would subsequently enable better resource allocation for mitigation and reduce human-wildlife conflicts (Lambertucci et al. 2015, Werber 2024).

Finally, using the morphotype approach together with complementary techniques (temporal, behavioral and spatial variability quantification), we can start discussing “Aerodiversity” as a measure of system health and conservation significance, like in terrestrial or aquatic ecosystems (Brooks et al. 2006, Maclaurin and Sterelny 2008). More finely segregated radar datasets can also improve predictions of abundance and behavior of flying animals, allowing researchers to test ecological theories in the aerial habitats (Bloch et al. 2025, Nilsson et al. 2025), more effectively manage human activity in the airspace, and better forecast aerial animals’ responses to large-scale processes like climate change.

## Acknowledgments

I thank Nir Sapir, Jason Chapman and Michael Kaluzhny for their valuable advice throughout. I further thank Yoav Perlman for support in the taxonomic analysis. Financial support to this study was provided by the Israeli Science Foundation (Award numbers 1652/22 and 2333/17) and the University of Haifa. I wish to thank the Hula Research center team for providing infrastructure and housing the radar station used in the taxonomic analysis.

