## Supplementary material for "The morphotype approach to classification of aerial animals in radar data": Code: Aerial_morphytype_script.Rmd

````
---
title: "preparation of hourly summary table"
author: "Yuval Werber"
date: "2022-10-19"
output: html_document
---

---
title: "Muiltivariate_analysis_israel_radar"
author: "Yuval Werber"
date: "2022-09-15"
output: html_document
---

```{r setup, include=FALSE}
knitr::opts_chunk$set(echo = TRUE)
knitr::opts_chunk$set(root.dir="/Users/yuvalwerber/Work/Academy/Morphotype_exploration")
setwd("/Users/yuvalwerber/Work/Academy/Morphotype_exploration")
library(tidyr)
library(pacman)
p_load(ggplot2, dplyr, tidyverse, rmarkdown, mvabund,  gridExtra, MuMIn, visreg, car, lme4,lmerTest,lme4,pivottabler, data.table,blmeco, AER)
library(tidyverse)
library(rstatix)
library(ggpubr)


```


Load RADAR data

```{r}
getwd()
setwd("/Users/yuvalwerber/Work/Academy/Morphotype_exploration")
load("/Users/yuvalwerber/Work/Academy/Morphotype_exploration/Full_radar_data_270122") 

```

data processing

```{r}
#remove signals without wff
Full_radar_data_250122$WFF1<-as.numeric(Full_radar_data_250122$WFF1)
data<-Full_radar_data_250122%>%drop_na(WFF1)
rm(Full_radar_data_250122)
#Keep Hula information
data<-data%>%filter(Location=="Hula")

##Remove non-bird echoes
unique(data$predicted_class)
##insects
data<-data%>%filter(predicted_class!="insect")
##Flocks
data<-data%>%filter(predicted_class!="flock")
##other
data<-data%>%filter(predicted_class!="other")
##aeroplane
data<-data%>%filter(predicted_class!="aeroplane")
##bats
load("/Users/yuvalwerber/Work/Academy/Morphotype_exploration/Classified_radar_data_full_nospeed_bats")

c<-ifelse(data$ordinal_radar%in%Classified_radar_data_full_nospeed_bats$ordinal_radar,"Bat",data$predicted_class)
table(c)
data$predicted_class<-c
data<-data%>%filter(predicted_class!="Bat")


#Keep short pulse

data$Pulsetype<-as.factor(data$Pulsetype)
data<-data%>%filter(Pulsetype!="Long")

##remove detection altitude<50, detection altitude >1000
data$Altitude_m_AGL<-as.numeric(data$Altitude_m_AGL)
data<-data%>%filter(Altitude_m_AGL<1000)
data<-data%>%filter(Altitude_m_AGL>50)


#keep high quality classification by the classification p values
#(Scmihdt says >0.5)


#Passerines
data$Passerine_p<-as.numeric(data$Passerine_p)
passerines<-data[(data$predicted_class=="passerine" & data$Passerine_p>=50),]
hist(passerines$Passerine_p)

##Waders
data$Wader_p<-as.numeric(data$Wader_p)
Waders<-data[(data$predicted_class=="wader"& data$Wader_p>=50),]
hist(Waders$Wader_p)

##Bird
data$Bird_p<-as.numeric(data$Bird_p)
Birds<-data[(data$predicted_class=="bird" & data$Bird_p>=50),]
hist(Birds$Bird_p)

###LSB
data$LSB_p<-as.numeric(data$LSB_p)
LSBs<-data[(data$predicted_class=="Large single bird" & data$LSB_p>=50),]
hist(LSBs$LSB_p)

data_1<-rbind(Birds,LSBs,passerines,Waders)
data_1$Full_Class<-data_1$predicted_class
rm(Birds,passerines,Waders,LSBs,Classified_radar_data_full_nospeed_bats,data,c)
```

For each class, we will now conduct multiple comparsions for diameter over wff intervals based on tokey and lm (aov), to decide how to divide groups of classes by WFF. where the difference between to intervals is significant, we weill set a break.
###Chevk increase in 0.95 quantile of rcs for each group in 1 WFF intervals

```{r}
#Keep relevant parameters
RCS_test_data<-data_1[,c("predicted_class","WFF1","RCS(M^2)","SQRT_RCS(M)","Day_night", "Polarisation_ratio", "Absolute_polarisation", "Maximum_level")]
##Make WFF intervals:
RCS_test_data<-RCS_test_data%>%drop_na(WFF1)

#cut by 1 or 2 hz interval to check what works for each class
RCS_test_data$WFF_round<-round(RCS_test_data$WFF1,digits = 0)
RCS_test_data$WFF_cut2<-cut(RCS_test_data$WFF_round, seq(from = min(RCS_test_data$WFF_round), to = max(RCS_test_data$WFF_round), by = 2 ), labels=FALSE)
RCS_test_data$WFF_cut2label<-cut(RCS_test_data$WFF_round, seq(from = min(RCS_test_data$WFF_round), to = max(RCS_test_data$WFF_round), by = 2 ))

RCS_test_data$WFF_cut1<-cut(RCS_test_data$WFF_round, seq(from = min(RCS_test_data$WFF_round), to = max(RCS_test_data$WFF_round), by = 1 ), labels=FALSE)
RCS_test_data$WFF_cut1label<-cut(RCS_test_data$WFF_round, seq(from = min(RCS_test_data$WFF_round), to = max(RCS_test_data$WFF_round), by = 1 ))


##Calculate object diameter, a proxy of head,body length in birds
RCS_test_data$`RCS(M^2)`<-as.numeric(RCS_test_data$`RCS(M^2)`)
RCS_test_data$`SQRT_RCS(M)`<-as.numeric(RCS_test_data$`SQRT_RCS(M)`)
RCS_test_data$Diameter<-200*sqrt(RCS_test_data$`RCS(M^2)`/pi)

RCS_test_data<-RCS_test_data%>%drop_na(`RCS(M^2)`)
RCS_test_data<-RCS_test_data%>%drop_na(`SQRT_RCS(M)`)


##make the summary table with 0.9 and 0.95 quenatiles


q<-c(.9,.95)
options(scipen=999)

Summary_table<-RCS_test_data %>%
  group_by(predicted_class, WFF_cut1label) %>%
  summarize(n=n(),
    quant90rcs = quantile(`RCS(M^2)`, probs = q[1]), 
            quant95rcs = quantile(`RCS(M^2)`, probs = q[2]),
    quant90sqrt = quantile(`SQRT_RCS(M)`, probs = q[1]), 
            quant95sqrt = quantile(`SQRT_RCS(M)`, probs = q[2]),
    quant90sqrt = quantile(Diameter, probs = q[1]), 
            quant95sqrt = quantile(Diameter, probs = q[2]))


write.csv(Summary_table,"RCS_and_Diameter_pecerntiles_WFF_inteval_1.csv", row.names = FALSE)
getwd()

```
https://www.r-bloggers.com/2019/01/quantile-regression-in-r-2/
Now we want to know which wff cuts are separapble for each group

Extract upper 95 qunatile for each group=class*WFF interval by 1

```{r}

RCS_test_data$group<-paste(RCS_test_data$predicted_class,RCS_test_data$WFF_cut1)

#filtering according to group and quntile
library(dplyr)
RCS_test_data_95_cut_1<-RCS_test_data %>%
   group_by(group) %>%
   filter(`RCS(M^2)` > quantile(`RCS(M^2)`, probs = 0.95))

##make data with 2 hz intervals as well

RCS_test_data$group<-paste(RCS_test_data$predicted_class,RCS_test_data$WFF_cut2)

RCS_test_data_95_cut_2<-RCS_test_data %>%
   group_by(group) %>%
   filter(`RCS(M^2)` > quantile(`RCS(M^2)`, probs = 0.95))

```


```{r}

#Check wich groups are to small (arbitrarily smaller than 30, maybe this will change later on

RCS_test_data_95_cut_1<- RCS_test_data_95_cut_1 %>% 
  group_by(group) %>% 
  filter(n() >= 30)


RCS_test_data_95_cut_2<- RCS_test_data_95_cut_2 %>% 
  group_by(group) %>% 
  filter(n() >= 30)

```

Multiple comparison procedure for each radar predicted class
Passerines
```{r}
#start with 1 hz breaks

Passerines<-RCS_test_data_95_cut_1%>%filter(predicted_class=="passerine")


ggplot(Passerines, aes(x=as.factor(WFF_cut1label), y=Diameter))+ylim(0,30)+
    geom_boxplot()+facet_grid(rows =vars(Day_night))
##Day and night are very similar
 Danova<-aov((Passerines$Diameter~Passerines$Day_night ));Danova
  summary(Danova) 
  


Passerines$WFF_cut1<-as.factor(Passerines$WFF_cut1)
 Danova<-aov((Passerines$Diameter~Passerines$WFF_cut1 ));Danova
  summary(Danova) 
  
  TukeyHSD(Danova)


  unique(Passerines$WFF_cut1label)
  #Combining the wff intervals basd on the tokey comprisons
Passerines$Radar_class<-if_else(Passerines$WFF_round<8,1,
           if_else(Passerines$WFF_round<9,2,
                 if_else(Passerines$WFF_round<12,3,
                         if_else(Passerines$WFF_round<13,4,
                                 if_else(Passerines$WFF_round<14,5,
                                         if_else(Passerines$WFF_round<15,6,
                                                 if_else(Passerines$WFF_round<16,7,
                                                         if_else(Passerines$WFF_round<17,8,
                                                                 if_else(Passerines$WFF_round<18,9,
                                                                         if_else(Passerines$WFF_round<19,10,11))))))))))
unique(Passerines$Radar_class)
##break data into morphytype accordign to where difference between adjacent WFF intervals was signficant
Passerine_morphytypes<-Passerines %>%
  group_by(Radar_class) %>%
  summarize(n=n(),
  Mean_WFF=mean(WFF1),
  Mean_diameter=mean(Diameter),
  Mean_RCS=mean(`RCS(M^2)`)
  )

```


Waders
```{r}
#start with 1 hz breaks

Waders<-RCS_test_data_95_cut_1%>%filter(predicted_class=="wader")


ggplot(Waders, aes(x=as.factor(WFF_cut1), y=Diameter))+ylim(0,30)+
    geom_boxplot()+facet_grid(rows =vars(Day_night))


Waders$WFF_cut1<-as.factor(Waders$WFF_cut1)
 Danova<-aov((Waders$Diameter~Waders$WFF_cut1 ));Danova
  summary(Danova) 

  TukeyHSD(Danova)
  
  range(Waders$WFF1)
   unique(Passerines$WFF_cut1label)
  #Combining the wff intervals basd on the tokey comprisons
   Waders$WFF_cut1<-as.numeric(Waders$WFF_cut1)
unique(Waders$WFF_cut1)
   Waders$Radar_class<-if_else(Waders$WFF_cut1<3,1,
           if_else(Waders$WFF_cut1<4,2,
                 if_else(Waders$WFF_cut1<6,3,
                         if_else(Waders$WFF_cut1<7,4,
                                 if_else(Waders$WFF_cut1<8,5,
                                         if_else(Waders$WFF_cut1<9,6,
                                                 if_else(Waders$WFF_cut1<13,7,8)))))))
unique(Waders$Radar_class)
##break data into morphytype accordign to where difference between adjacent WFF intervals was signficant
Wader_morphytypes<-Waders %>%
  group_by(Radar_class) %>%
  summarize(n=n(),
  Mean_WFF=mean(WFF1),
  Mean_diameter=mean(Diameter),
  Mean_RCS=mean(`RCS(M^2)`)
  )

```


lsbs used WFFbreak 2 here
```{r}
#start with 1 hz breaks

lsbs<-RCS_test_data_95_cut_1%>%filter(predicted_class=="Large single bird")

lsbs$WFF_cut2<-as.factor(lsbs$WFF_cut2)
 Danova<-aov((lsbs$Diameter~lsbs$WFF_cut2 ));Danova
  summary(Danova) 
    TukeyHSD(Danova)
  
  #Combining the wff intervals basd on the tokey comprisons
   lsbs$WFF_cut2<-as.numeric(lsbs$WFF_cut2)
unique(lsbs$WFF_cut2)
   lsbs$Radar_class<-if_else(lsbs$WFF_cut2<2,1,2)
   
   range(lsbs$WFF_cut2)
ggplot(lsbs, aes(x=as.factor(WFF_cut2), y=Diameter))+
    geom_boxplot()  
  

LSB_morphytypes<-lsbs %>%
  group_by(Radar_class) %>%
  summarize(n=n(),
  Mean_WFF=mean(WFF1),
  Mean_diameter=mean(Diameter),
  Mean_RCS=mean(`RCS(M^2)`)
  )

```

bird
```{r}

birds<-RCS_test_data_95_cut_1%>%filter(predicted_class=="bird")

ggplot(birds, aes(x=as.factor(WFF_cut1), y=Diameter))+
    geom_boxplot()+facet_grid(rows =vars(Day_night))

birds$WFF_cut1<-as.factor(birds$WFF_cut1)
 Danova<-aov((birds$Diameter~birds$WFF_cut1 ));Danova
  summary(Danova) 
options(max.print = 2000)  
  TukeyHSD(Danova)
  
birds$WFF_cut1<-as.numeric(birds$WFF_cut1)
     birds$Radar_class<-if_else(birds$WFF_cut1<2,1,
           if_else(birds$WFF_cut1<3,2,
                 if_else(birds$WFF_cut1<4,3,
                         if_else(birds$WFF_cut1<5,4,
                                 if_else(birds$WFF_cut1<8,5,
                                         if_else(birds$WFF_cut1<9,6,
                                                 if_else(birds$WFF_cut1<11,7,
                                                         if_else(birds$WFF_cut1<16,8,
                                                                 if_else(birds$WFF_cut1<17,9,10)))))))))
unique(birds$Radar_class)
  

ggplot(birds, aes(x=as.factor(WFF_cut1), y=Diameter))+
    geom_boxplot() 

ggplot(birds, aes(x=as.factor(Radar_class), y=Diameter))+
    geom_boxplot() 
  
##Difference between 12-13 and 18-19 only. looks like the second comparsion in linn is usually 
  ##Good differences in cut 1
  ##no need for cut 2
Birds_morphytypes<-birds %>%
  group_by(Radar_class) %>%
  summarize(n=n(),
  Mean_WFF=mean(WFF1),
  Mean_diameter=mean(Diameter),
  Mean_RCS=mean(`RCS(M^2)`)
  )
```


Boxplots for paper 
Figure 1
```{r}
unique(RCS_test_data_95_cut_1$predicted_class)
RCS_test_data_95_cut_1$predicted_class<-ifelse(RCS_test_data_95_cut_1$predicted_class=="passerine","Passerine",ifelse(RCS_test_data_95_cut_1$predicted_class=="wader","Wader",ifelse(RCS_test_data_95_cut_1$predicted_class=="bird","Bird","Large single bird")))

RCS_test_data_95_cut_1$predicted_class<-factor(RCS_test_data_95_cut_1$predicted_class,levels =c("Large single bird","Passerine","Wader","Bird") )

boxplot<-ggplot(RCS_test_data_95_cut_1, aes(x=as.factor(WFF_cut1), y=Diameter))+
    geom_boxplot(outlier.size = 0.5,outlier.color = "grey")+facet_wrap(.~predicted_class,ncol=1,scales="free")+theme_bw()+ylab("Body-head length (cm)")+xlab("Wing flapping frequency (Hz)")+theme(strip.text = element_text(size = 14))+theme(
    strip.background = element_rect(fill = "white"),strip.text = element_text(face = "bold"))


##adding morphytype breaks

plot<-ggdraw() +
  draw_plot(boxplot, x = 0, y = 0, width =1 , height = 1) +
  draw_line(x = c(0.62, 0.62), y = c(0.775, 0.961), color = "darkred", size = 1.)+#LSB_1
   draw_line(x = c(0.195, 0.195), y = c(0.532, 0.718), color = "darkred", size = 1.)+#Pass_1
     draw_line(x = c(0.251, 0.251), y = c(0.532, 0.718), color = "darkred", size = 1.)+#Pass_2
     draw_line(x = c(0.42, 0.42), y = c(0.532, 0.718), color = "darkred", size = 1.)+#Pass_3
   draw_line(x = c(0.4765, 0.4765), y = c(0.532, 0.718), color = "darkred", size = 1.)+#Pass_4
   draw_line(x = c(0.533, 0.533), y = c(0.532, 0.718), color = "darkred", size = 1.)+#Pass_5
   draw_line(x = c(0.5895, 0.5895), y = c(0.532, 0.718), color = "darkred", size = 1.)+#Pass_6
   draw_line(x = c(0.646, 0.646), y = c(0.532, 0.718), color = "darkred", size = 1.)+#Pass_7
   draw_line(x = c(0.7025, 0.7025), y = c(0.532, 0.718), color = "darkred", size = 1.)+#Pass_8
   draw_line(x = c(0.7590, 0.7590), y = c(0.532, 0.718), color = "darkred", size = 1.)+#Pass_9
   draw_line(x = c(0.8155, 0.8155), y = c(0.532, 0.718), color = "darkred", size = 1.)+#Pass_10
   draw_line(x = c(0.1235, 0.1235), y = c(0.288, 0.475), color = "darkred", size = 1.)+#Wader_1
   draw_line(x = c(0.167, 0.167), y = c(0.288, 0.475), color = "darkred", size = 1.)+#Wader_2
  draw_line(x = c(0.254, 0.254), y = c(0.288, 0.475), color = "darkred", size = 1.)+#Wader_3
  draw_line(x = c(0.296, 0.296), y = c(0.288, 0.475), color = "darkred", size = 1.)+#Wader_4
  draw_line(x = c(0.339, 0.339), y = c(0.288, 0.475), color = "darkred", size = 1.)+#Wader_5
  draw_line(x = c(0.382, 0.382), y = c(0.288, 0.475), color = "darkred", size = 1.)+#Wader_6
  draw_line(x = c(0.554, 0.554), y = c(0.288, 0.475), color = "darkred", size = 1.)+#Wader_7
    draw_line(x = c(0.1131, 0.1131), y = c(0.044, 0.231), color = "darkred", size = 1.)+#Bird_1
  draw_line(x = c(0.1466, 0.1466), y = c(0.044, 0.231), color = "darkred", size = 1.)+#Bird_2
  draw_line(x = c(0.1801, 0.1801), y = c(0.044, 0.231), color = "darkred", size = 1.)+#Bird_3
  draw_line(x = c(0.2136, 0.2136), y = c(0.044, 0.231), color = "darkred", size = 1.)+#Bird_4
  draw_line(x = c(0.3145, 0.3145), y = c(0.044, 0.231), color = "darkred", size = 1.)+#Bird_5
  draw_line(x = c(0.348, 0.348), y = c(0.044, 0.231), color = "darkred", size = 1.)+#Bird_6
  draw_line(x = c(0.4150, 0.4150), y = c(0.044, 0.231), color = "darkred", size = 1.)+#Bird_7
  draw_line(x = c(0.583, 0.583), y = c(0.044, 0.231), color = "darkred", size = 1.)+#Bird_8
   draw_line(x = c(0.6166, 0.6166), y = c(0.044, 0.231), color = "darkred", size = 1.)+
     draw_label("L1",x = 0.375,y = 0.95,hjust = 0.5,vjust = 0.5,fontface = "bold",color = "darkred",size = 12)+
    draw_label("L2",x = 0.8,y = 0.95,hjust = 0.5,vjust = 0.5,fontface = "bold",color = "darkred",size = 12)+
    draw_label("P1",x = 0.14,y = 0.71,hjust = 0.5,vjust = 0.5,fontface = "bold",color = "darkred",size = 12)+
  draw_label("P2",x = 0.222,y = 0.71,hjust = 0.5,vjust = 0.5,fontface = "bold",color = "darkred",size = 12)+
  draw_label("P3",x = 0.333,y = 0.71,hjust = 0.5,vjust = 0.5,fontface = "bold",color = "darkred",size = 12)+
  draw_label("P4",x = 0.449,y = 0.71,hjust = 0.5,vjust = 0.5,fontface = "bold",color = "darkred",size = 12)+
  draw_label("P5",x = 0.505,y = 0.71,hjust = 0.5,vjust = 0.5,fontface = "bold",color = "darkred",size = 12)+
  draw_label("P6",x = 0.561,y = 0.71,hjust = 0.5,vjust = 0.5,fontface = "bold",color = "darkred",size = 12)+
  draw_label("P7",x = 0.617,y = 0.71,hjust = 0.5,vjust = 0.5,fontface = "bold",color = "darkred",size = 12)+
  draw_label("P8",x = 0.673,y = 0.71,hjust = 0.5,vjust = 0.5,fontface = "bold",color = "darkred",size = 12)+
  draw_label("P9",x = 0.729,y = 0.71,hjust = 0.5,vjust = 0.5,fontface = "bold",color = "darkred",size = 12)+
  draw_label("P10",x = 0.786,y = 0.71,hjust = 0.5,vjust = 0.5,fontface = "bold",color = "darkred",size = 12)+
  draw_label("P11",x = 0.9,y = 0.71,hjust = 0.5,vjust = 0.5,fontface = "bold",color = "darkred",size = 12)+
   draw_label("W1",x = 0.099,y = 0.465,hjust = 0.5,vjust = 0.5,fontface = "bold",color = "darkred",size = 12)+
  draw_label("W2",x = 0.146,y = 0.465,hjust = 0.5,vjust = 0.5,fontface = "bold",color = "darkred",size = 12)+
  draw_label("W3",x = 0.21,y = 0.465,hjust = 0.5,vjust = 0.5,fontface = "bold",color = "darkred",size = 12)+
  draw_label("W4",x = 0.275,y = 0.465,hjust = 0.5,vjust = 0.5,fontface = "bold",color = "darkred",size = 12)+
  draw_label("W5",x = 0.317,y = 0.465,hjust = 0.5,vjust = 0.5,fontface = "bold",color = "darkred",size = 12)+
  draw_label("W6",x = 0.361,y = 0.465,hjust = 0.5,vjust = 0.5,fontface = "bold",color = "darkred",size = 12)+
  draw_label("W7",x = 0.471,y = 0.465,hjust = 0.5,vjust = 0.5,fontface = "bold",color = "darkred",size = 12)+
  draw_label("W8",x = 0.773,y = 0.465,hjust = 0.5,vjust = 0.5,fontface = "bold",color = "darkred",size = 12)+
  draw_label("B1",x = 0.095,y = 0.222,hjust = 0.5,vjust = 0.5,fontface = "bold",color = "darkred",size = 12)+
  draw_label("B2",x = 0.129,y = 0.222,hjust = 0.5,vjust = 0.5,fontface = "bold",color = "darkred",size = 12)+
  draw_label("B3",x = 0.163,y = 0.222,hjust = 0.5,vjust = 0.5,fontface = "bold",color = "darkred",size = 12)+
  draw_label("B4",x = 0.197,y = 0.222,hjust = 0.5,vjust = 0.5,fontface = "bold",color = "darkred",size = 12)+
  draw_label("B5",x = 0.262,y = 0.222,hjust = 0.5,vjust = 0.5,fontface = "bold",color = "darkred",size = 12)+
  draw_label("B6",x = 0.331,y = 0.222,hjust = 0.5,vjust = 0.5,fontface = "bold",color = "darkred",size = 12)+
  draw_label("B7",x = 0.381,y = 0.222,hjust = 0.5,vjust = 0.5,fontface = "bold",color = "darkred",size = 12)+
  draw_label("B8",x = 0.501,y = 0.222,hjust = 0.5,vjust = 0.5,fontface = "bold",color = "darkred",size = 12)+
   draw_label("B9",x = 0.599,y = 0.222,hjust = 0.5,vjust = 0.5,fontface = "bold",color = "darkred",size = 12)+
  draw_label("B10",x = 0.83,y = 0.222,hjust = 0.5,vjust = 0.5,fontface = "bold",color = "darkred",size = 12)
  
  
  
  
  
   
  


ggsave("WFF_BOXPLOTS11.png", width =7 , height =10,dpi=300)
getwd()
```
Morphotype prevalence analisys

```{r}

Morphotype_prevalence<-read.csv("Israel_bird_Species_no_desert.csv")
Monthly_prevalence<-read.csv("Month_morphotype_prevalence.csv")


##Start with morphytype hustogram

Morphotype<-Morphotype_prevalence%>%
  group_by(Radar.class,Morphtype)%>%
  dplyr::summarise(Species=n()
   #V_test_vector_length=v_test_statistic(r_test_data),
   #V_test_p_value = v_test_p(r_test_data)
      )

hist(Morphotype$Species)

max(Morphotype$Species)

Monthly<-Monthly_prevalence%>%
  group_by(Morphotype_month)%>%
  dplyr::summarise(Species=n()
   #V_test_vector_length=v_test_statistic(r_test_data),
   #V_test_p_value = v_test_p(r_test_data)
      )

Monthly_summary<-Monthly%>%
  group_by(Species)%>%
  dplyr::summarise(count=n()
   #V_test_vector_length=v_test_statistic(r_test_data),
   #V_test_p_value = v_test_p(r_test_data)
      )

hist(Monthly$Species)


# Create the histogram
  Monthly_hist<-ggplot(Monthly, aes(x = Species)) +
  geom_histogram(color="black",fill="lightgrey",bins = 50)+theme_light()+xlab("Number of species in Morphytype*Month")+ylab("Number of Morphotype*Months")+scale_x_continuous(breaks = seq(0, 50, by = 2))+
  theme(axis.text.x = element_text(angle = 45, hjust = 1))+scale_y_continuous(breaks = seq(0, 70, by = 5))+
  geom_label(aes(x=26, y=66.3, label="Morphotype.Months"), fill = "white",fontface = "bold",size = 6)
  
   Morphotype_hist<- ggplot(Morphotype, aes(x = Species)) +
  geom_histogram(color="black",fill="lightgrey",bins = 50)+theme_light()+xlab("Number of species in Morphytype")+ylab("Number of Morphotypes")+scale_x_continuous(breaks = seq(0, 54, by = 2))+
  theme(axis.text.x = element_text(angle = 45, hjust = 1))+
  geom_label(aes(x=27, y=4.73, label="Morphotypes"), fill = "white",fontface = "bold",size = 6)
   
  
   plot<-ggdraw() +
  draw_plot(Morphotype_hist, x = 0, y = 0.5, width =1, height = 0.5) +
  draw_plot(Monthly_hist, x = 0, y = 0, width =1, height = 0.5) +
  draw_label("A.",x = 0.03,y = 0.98,hjust = 0.5,vjust = 0.5,fontface = "bold",size = 12)+
  draw_label("B.",x = 0.03,y = 0.48,hjust = 0.5,vjust = 0.5,fontface = "bold",size = 12)
ggsave("Figure2.png", width = 5, height = 8,dpi=300)   
write.csv(Morphotype_prevalence,file= "Table S2.csv")
```

Morphotype description table
```{r}


Table<-rbind( Passerine_morphytypes,Birds_morphytypes,Wader_morphytypes,LSB_morphytypes )

write_csv(Table, file = "Morphotype_table.csv")
```
````
